# Shared and divergent transcriptomic regulation in nucleus accumbens D1 and D2 medium spiny neurons by cocaine and morphine

**DOI:** 10.1101/2023.09.19.558477

**Authors:** Caleb J Browne, Philipp Mews, Xianxiao Zhou, Leanne M Holt, Molly Estill, Rita Futamura, Anne Schaefer, Paul J Kenny, Yasmin L Hurd, Li Shen, Bin Zhang, Eric J Nestler

## Abstract

Substance use disorders (SUDs) induce widespread molecular dysregulation in the nucleus accumbens (NAc), a brain region pivotal for coordinating motivation and reward. These molecular changes are thought to support lasting neural and behavioral disturbances that promote drug-seeking in addiction. However, different drug classes exert unique influences on neural circuits, cell types, physiology, and gene expression despite the overlapping symptomatology of SUDs. To better understand common and divergent molecular mechanisms governing SUD pathology, our goal was to survey cell-type-specific restructuring of the NAc transcriptional landscape in after psychostimulant or opioid exposure. We combined fluorescence-activated nuclei sorting and RNA sequencing to profile NAc D1 and D2 medium spiny neurons (MSNs) across cocaine and morphine exposure paradigms, including initial exposure, prolonged withdrawal after repeated exposure, and re-exposure post-withdrawal. Our analyses reveal that D1 MSNs display many convergent transcriptional responses across drug classes during exposure, whereas D2 MSNs manifest mostly divergent responses between cocaine and morphine, with morphine causing more adaptations in this cell type. Utilizing multiscale embedded gene co-expression network analysis (MEGENA), we discerned transcriptional regulatory networks subserving biological functions shared between cocaine and morphine. We observed largely integrative engagement of overlapping gene networks across drug classes in D1 MSNs, but opposite regulation of key D2 networks, highlighting potential therapeutic gene network targets within MSNs. These studies establish a landmark, cell-type-specific atlas of transcriptional regulation induced by cocaine and by morphine that can serve as a foundation for future studies towards mechanistic understanding of SUDs. Our findings, and future work leveraging this dataset, will pave the way for the development of targeted therapeutic interventions, addressing the urgent need for more effective treatments for cocaine use disorder and enhancing the existing strategies for opioid use disorder.

## Introduction

Substance use disorders (SUDs) represent a spectrum of chronic, relapsing neuropsychiatric conditions marked by an individual’s persistent compulsion to seek and consume drugs at the expense of healthy goals. Drugs of abuse are thought to gain their ability to control motivated behavior by causing persistent functional and molecular restructuring of the brain reward circuit, ultimately tuning behavior towards drug-seeking and drug-taking (1–3). Drug-induced molecular changes also persist long after drug use is terminated, and this is thought to support lasting changes to reward circuit function that can drive relapse – one of the most challenging aspects of SUD treatment (4). Thus, understanding the mechanisms governing persistent drug-induced molecular changes may hold the key to developing effective therapeutic interventions that can prevent relapse and promote sustained recovery.

Cocaine use disorder (CUD) and Opioid use disorder (OUD) are two prominent forms of SUDs that have garnered significant attention due to their widespread prevalence and societal impact. Although both are considered a form of SUD, the manifestation of CUD and OUD are different; cocaine and opioids exert distinct pharmacological effects, elicit overlapping but also some unique physiological changes to brain reward circuits (5), and may induce separable patterns of molecular remodeling supporting long-term disease pathogenesis (6). Further, withdrawal and abstinence in CUD are less severe than in OUD, and long-term abstinence without pharmacotherapeutic intervention is nearly impossible in OUD (7,8). Central to the neurobiology of both SUDs is the nucleus accumbens (NAc), a critical region in the brain’s reward circuitry, which undergoes significant transcriptional alterations in response to chronic drug exposure. Recent studies have begun to elucidate transcriptional disruptions in the NAc associated with CUD and OUD (9–12) revealing both overlapping and divergent molecular signatures. For instance, both disorders have been linked to alterations in genes associated with biological processes supporting synaptic plasticity, neurotransmission, and intracellular signaling. However, the specific genes and pathways affected, as well as the magnitude and direction of these changes, can vary between CUD and OUD.

A deeper understanding of the molecular underpinnings of SUDs necessitates a cell-type-specific perspective, given the heterogeneous nature of the NAc. Two primary medium spiny neuron (MSN) subtypes, distinguished by their dopamine receptor expression—D1- and D2-MSNs—dominate the NAc. These MSN subtypes exhibit unique input-output architecture, play distinct roles in goal-directed behavior, and their dysregulation is thought to contribute differently to the pathophysiology of SUDs (13–16). Recent investigations into CUD and OUD mouse models have highlighted differential transcriptional responses in D1 and D2 MSNs (17,18). These studies suggest that D1 MSNs primarily undergo transcriptional activation in response to drug exposure, while D2 MSNs might exhibit more complex patterns of gene regulation (19). Despite these advances, we lack a complete picture of how the transcriptional landscape defining D1 and D2 MSN function evolves across different stages of the addiction cycle for either cocaine or opioids. Investigating how common vs. distinct molecular pathways in these two cell types are dynamically reprogrammed across CUD and OUD development and maintenance will be crucial for identifying key gene networks that may serve as potential therapeutic targets.

To address this knowledge gap, we explored transcriptome-wide disruptions induced by cocaine or morphine in D1 and D2 MSNs. We examined the molecular changes both acutely and following prolonged withdrawal from chronic exposure. Furthermore, we sought to understand how re-exposure to these drugs after a withdrawal period compares to the acute response. This design allowed us to capture the transcriptional shifts across various drug exposure and withdrawal stages, providing a comprehensive view of the molecular adaptations in D1 and D2 MSNs in response to cocaine and to morphine. Currently, there are no FDA-approved treatments for CUD (20), and while there are treatments available for OUD they remain lacking in efficacy and accessibility (7,8). Addressing these therapeutic inadequacies is of paramount importance. Our findings provide new insight into the long-lasting and MSN subtype-specific dysregulation of transcription triggered in the NAc by cocaine and by morphine, laying the groundwork for new avenues toward therapeutic development and potential interventions for people experiencing SUDs.

## Results

Our goal was to create and mine a cell-type-specific atlas of molecular regulation in the NAc induced by cocaine or morphine across distinct addiction-relevant drug exposure paradigms (Fig. 1A). To specifically assay changes in D1 or D2 MSN transcriptional regulation, we crossed D1-Cre or D2-Cre mice with EGFP-L10a mice, which express a Cre-dependent ribosome-tagged GFP reporter, enabling cell-type-specific nuclei purification. Mice were treated with cocaine (20 mg/kg) or morphine (10 mg/kg), at doses known to cause strong rewarding effects in mice (21), or with saline, for 10-15 days, followed by 30 days of long-term withdrawal. The 30 d timepoint was chosen to coincide with a period of abstinence associated with high levels of drug craving that promotes relapse (22), and has been linked to dynamic transcriptional regulation within the NAc for both cocaine and opioids (9,11). At the 30 d time point, animals received a challenge injection of either saline, cocaine, or morphine in the homecage and were euthanized 1h later, at which point the NAc was dissected and flash frozen. This design produced three exposure conditions in which to examine molecular mechanisms of SUDs: acute, first-ever exposure (SC/SM), long-term withdrawal from chronic exposure (CS/MS), and re-exposure after protracted withdrawal (CC/MM). Nuclei from D1 or D2 NAc MSNs were then purified using FANS and processed for RNAseq analysis. 6-9 mice were analyzed per group across all conditions (see Supplementary Table 1 for details). Differential expression analysis comparing all drug exposure conditions with their respective control group (SS) was performed using DEseq2 with independent filtering disabled, using only the top 50% of genes in the genome based on normalized read counts.

**Fig 1.**
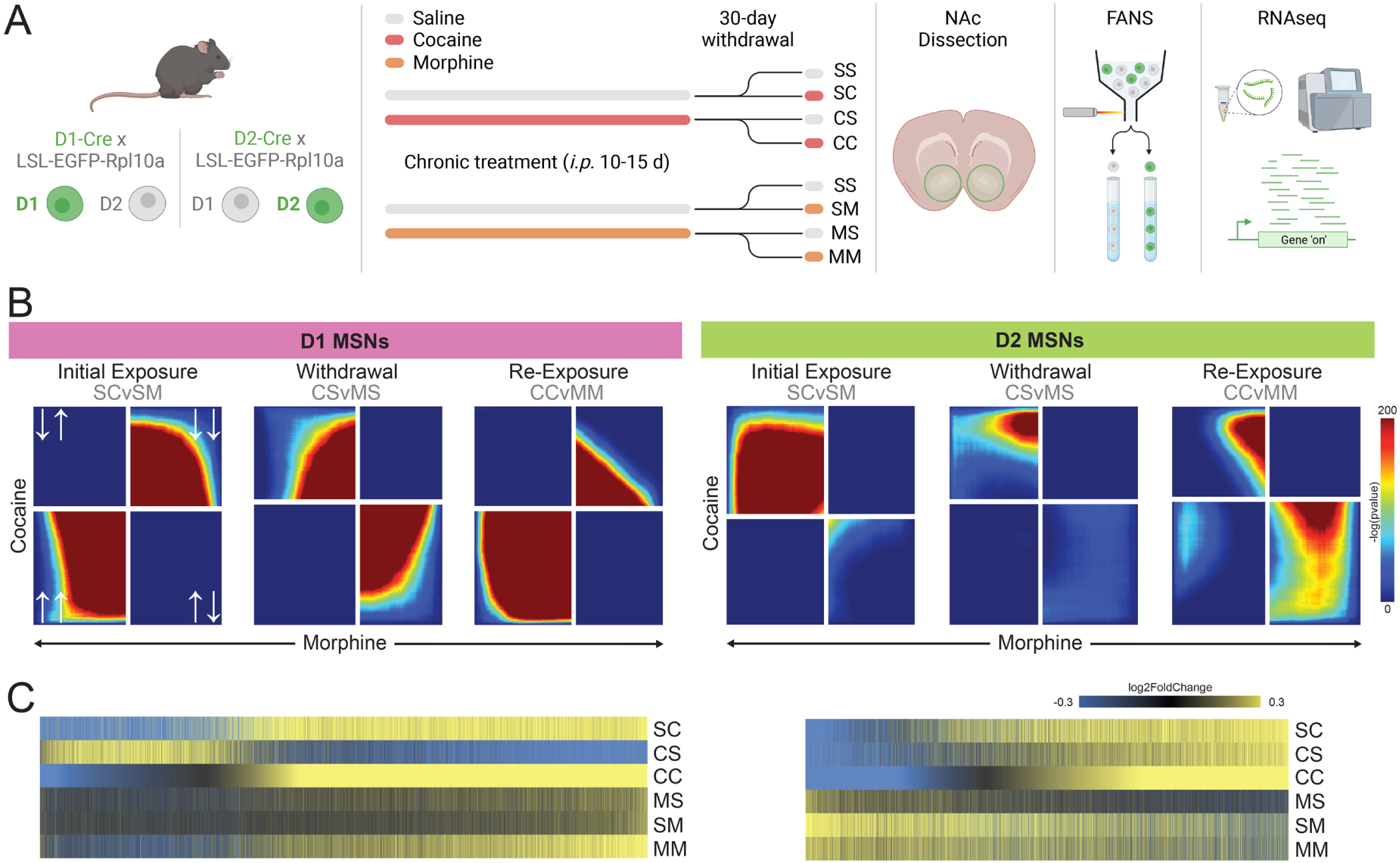
Patterns of transcriptional regulation in NAc D1 and D2 MSNs induced by cocaine or morphine. **A**, Experimental paradigm outlining cell-type-specific RNAseq after cocaine or morphine exposure conditions. Nuclei in D1 or D2 MSNs were GFP-tagged for purification by crossing mice with a floxed EGFP-L10a allele, which inserts a GFP tag into the ribosomal L10a locus, with D1-Cre or D2-Cre mice. Double-transgenic mice were treated with saline, cocaine (20 mg/kg), or morphine (10 mg/kg) for 10 days (cocaine cohort) or 15 days (morphine cohort), followed by a 30-day homecage withdrawal period. Subsequently, mice were challenged with an injection of either saline, cocaine, or morphine and were sacrificed 1h later from the homecage. NAc tissue was extracted, GFP-positive nuclei were purified via FANS, and RNAseq was performed on these D1 and D2 MSN nuclei. Note, cocaine and morphine cohorts were sequenced at different times. **B**, Rank-rank hypergeometric overlap plots illustrating threshold-free transcriptomic similarities between cocaine and morphine across first-ever exposure (SC vs. SM), withdrawal (CS vs. MS), or re-exposure after withdrawal (CC vs MM) separately for D1 or D2 MSNs. Heat indicates strength of overlap, and quadrants represent direction of gene expression (white arrows; lower-left quadrant, genes up in both; upper-right quadrant, genes down in both; upper left quadrant, genes down in cocaine but up in morphine; lower right quadrant: genes up in cocaine but down in morphine) **C**, Union heatmaps seeded to the CC condition showing log2FoldChange of genes identified as significantly (>20% fold change, p<0.05) upregulated (yellow) or downregulated (blue) in all experimental conditions.

### Cocaine and morphine cause distinct patterns of transcriptional regulation in D1 and D2 MSNs

We first aimed to characterize general signatures of transcriptional changes in response to cocaine vs. morphine across D1 and D2 MSNs. To do this, we used rank-rank hypergeometric overlap (RRHO), which enables comparisons of transcriptomic regulation across experimental conditions in a threshold-free manner. For D1 MSNs (Fig 1B), RRHO plots demonstrate that first-ever exposure to cocaine (SC) or morphine (SM) elicits a strong overlapping transcriptional response. This pattern is preserved upon re-exposure to cocaine (CC) or morphine (MM) after withdrawal, with a high degree of overlap in upregulated genes across CC and MM conditions. However, we noted that during cocaine or morphine withdrawal (CS or MS), transcriptional regulation is anticorrelated. This suggests that cocaine and morphine induce a typical pattern of molecular regulation in D1 MSNs when the drug is on board but exert separable effects through prolonged withdrawal. On the other hand, in D2 MSNs (Fig 1C), RRHO plots show very little concordance across any cocaine or morphine condition, except for genes upregulated by first-ever morphine (SM) being downregulated by first-ever cocaine (SC). These results suggest that D1 MSNs exhibit broadly similar transcriptional regulation across drug classes during exposure, regardless of prior history, while transcriptional regulation in D2 MSNs is largely disparate between drug classes.

Having identified overlapping and distinct patterns of gene regulation in D1 and D2 MSNs, we next examined whether these changes related to shifts in gene priming and desensitization across drug exposure conditions. We generated union heatmaps for D1 or D2 MSNs that display all protein-coding genes identified to be significantly regulated (20% Fold Change, p<0.05) in at least 1 experimental condition, and presented the corresponding fold change for each condition irrespective of significance (Figure 1C). These heatmaps were then seeded to the CC condition to organize results around chronic drug treatment and re-exposure after withdrawal. For D1 MSNs, we found that transcriptional responses to morphine and cocaine were essentially parallel, except for the CS condition (cocaine withdrawal), consistent with RRHO findings. First-ever cocaine (SC) exposure elicits a transcriptional response that is largely reactivated upon re-exposure after withdrawal (CC), particularly for upregulated genes. However, this pattern is specific to the drug being on board; cocaine withdrawal (CS) induced a largely inverted gene expression signature compared to either SC or CC conditions. In fact, many genes inhibited in SC are activated in CS but no longer inhibited with re-exposure, indicative of a group of genes showing normalization of responses. The prolonged suppression of numerous genes in cocaine withdrawal suggests that D1-relevant biological functions cycle with repeated exposure to cocaine and are disrupted long-term through withdrawal. The pattern of inverted gene expression in D1 MSNs after withdrawal was not observed for morphine, which exhibits consistent, parallel regulation across SM, MS, and MM conditions, albeit to a diminished foldchange compared to cocaine. Notably, the effects of morphine are weak at first (SM) but are much more robust with re-exposure after withdrawal (MM), indicative of a sensitizing effect of repeated morphine exposure in D1 MSNs. We also found that both CC and MM show a high degree of overlapping up- and down-regulated genes, supporting the notion that D1 MSNs support convergent influences of cocaine and morphine in addiction-relevant drug exposure, although this shifts dramatically in withdrawal.

In contrast to D1 MSNs, D2 MSNs exhibit a distinct transcriptional response to cocaine vs. morphine across all exposure conditions (Figure 1C), paralleling changes observed in RRHO analysis. Unlike in D1 MSNs, cocaine shows a largely conserved pattern of gene regulation across acute exposure, withdrawal, and re-exposure, and the same is observed across these conditions for morphine. However, genes most strongly upregulated by cocaine are downregulated by morphine, and vice versa. These threshold-imposed results are consistent with RRHOs for D2 MSNs, which showed little transcriptomic overlap or anticorrelations between cocaine and morphine conditions. We also noted that morphine withdrawal (MS) elicited more robust transcriptional responses compared to either first-ever exposure (SM) or re-exposure (MM), which was not observed in D1 MSNs. This suggests that long-term withdrawal from morphine primes gene expression in D2 MSNs, which is normalized by re-exposure to morphine.

Together, these findings show that, while cocaine and morphine drive similar transcriptional responses in D1 MSNs, particularly while the drug is on board, these drugs exert separable or even opposite influences on transcriptional regulation in D2 MSNs. These results imply a unique and persistent influence that these two drug classes have on broad biological processes within the NAc.

### Cocaine consolidates unique transcriptional regulation between D1 and D2 MSNs, while morphine accumulates common transcriptional regulation

We next examined how transcriptional regulation differed across cell types within each exposure condition (Fig. S1). RRHO plots show that first-ever cocaine exposure elicits broadly similar patterns of transcriptional regulation across D1 and D2 MSNs, particularly in upregulated genes (Fig. S1). Interestingly, studies using single nucleus RNAseq have also found an overlap in transcriptional regulation between D1 and D2 MSNs in response to acute cocaine (18). These findings were somewhat surprising due to the well-established opposite cellular responses to D1 and D2 receptor action, wherein dopamine binding to D1 receptors causes neuronal activation, while D2 receptors suppress neural responses. At the molecular level, acute cocaine has been shown to consistently activate many first-wave immediate early genes (IEGs), including several transcription factors, in the NAc such as *Arc*, *Fos*, and *Jun* (23). These IEGs are thought to initiate waves of broad transcriptional regulation and intracellular processes required for neuroplasticity. Interestingly, despite the observed overlap in transcriptional response across D1 and D2 MSNs to first-ever cocaine (SC), these IEGs (*Arc*, Log2FC=1.36, p<0.05; *Fos*, Log2FC=1.17, p<0.05; *Jun*, Log2FC=1.36, p<0.05) were only upregulated in D1 MSNs, and unaffected in D2 MSNs. Thus, the transcriptomic convergence observed in RRHO analyses may be independent of intracellular responses to dopamine action at D1 and D2 receptors. Nonetheless, the initial overlap in transcriptional responses to cocaine between D1 and D2 MSNs appears to consolidate separately between cell types with a history of exposure. RRHOs comparing D1 and D2 MSNs for the CC condition find much less transcriptional overlap. The CC condition appears to cause genes that are typically downregulated in D1 MSNs to be upregulated in D2 MSNs. Taken with results from Figure 1, these findings suggest that, while many genes in SC and CC overlap within D1 and D2 MSNs, the particular genes that are activated in the CC condition likely coordinate distinct processes within each cell type.

First-ever morphine exposure (SM), on the other hand, induces an entirely unique pattern of transcriptional regulation across D1 and D2 MSNs (Fig S1). Additionally, unlike cocaine re-exposure which caused a restructuring of transcriptional regulation across cell types, morphine re-exposure (MM) induced more overlap in genes regulated across D1 and D2 MSNs, particularly for genes upregulated across these cell types. Interestingly, transcriptional regulation elicited by withdrawal from either cocaine (CS) or morphine (MS) was mostly distinct between D1 and D2 MSNs.

Together, these findings indicate that increasing cocaine exposure consolidates transcriptional regulation between D1 and D2 MSNs in the NAc, while morphine treatment progressively tunes similar patterns of molecular regulation across cell types.

### Unique gene network regulation across cocaine and morphine exposures

Having identified broad changes to gene expression within D1 and D2 NAc MSNs induced by cocaine or morphine, we next determined whether persistent molecular regulation was organized into gene networks that may subserve unique biological functions. Using multiscale embedded gene co-expression network analysis (MEGENA; 24), we generated basal gene co-expression networks for D1 and D2 MSNs by collapsing all experimental conditions across each cell type. An advantage of MEGENA is the ability to organize gene networks hierarchically, by progressively refining more extensive networks into more granular ones that cluster around highly interconnected hub genes. Interestingly, we found that overall D2 network topology was less correlated than D1s, leading to a fragmentation of network organization (as illustrated in Figure 4A). The D1 network had 41,095 edges, with an average correlation coefficient of 0.96824 (SD = 0.019221), while the D2 network had 38,724 edges and an average correlation coefficient of 0.94304 (SD = 0.034624), which was significantly different (p < 2.2e-16). These findings are consistent with our prior work outlining nuclear D1 and D2 molecular profiles from naïve samples, demonstrating that D2 MSNs exhibit many more lowly expressed transcripts compared to D1 MSNs (25). Therefore, the basal network organization of D1 and D2 MSNs appears to be dramatically different, emphasizing their separable roles in biological processes and validating the importance of analyzing these cell types separately.

We next overlaid DEG lists onto MEGENA networks to determine how drug exposure drives coordinated changes across gene networks. To do this, we represented D1 and D2 gene network architecture as sunburst plots (Figure 2) that enable illustration of parent-child network organization, and shaded distinct networks, depicted by rungs of the plot, as enriched with upregulated (yellow) or downregulated (blue) genes (Log2FC >20%, pvalue <0.05). In D1 MSNs, first-ever cocaine exposure (SC) caused both up- and downregulation of many gene networks. Interestingly, these changes were completely lost during withdrawal from chronic cocaine. Considering the high number of DEGs whose expression was largely inverted compared to acute cocaine exposure, it was surprising to find that few, if any, gene networks were enriched with DEGs in the CS condition. This result suggests that, while many genes are affected through withdrawal, these changes do not engage gene networks as when cocaine is on board. Re-exposure to cocaine after withdrawal (CC) drove a transcriptional response marked by primarily upregulated gene networks. Interestingly, while some modules overlap with SC, close examination shows that many of the genes in these modules, while stemming from the same parent module, are largely distinct from those upregulated by acute cocaine exposure. Further, the network compartmentalization of downregulated genes induced in SC was lost in CC.

**Fig 2.**
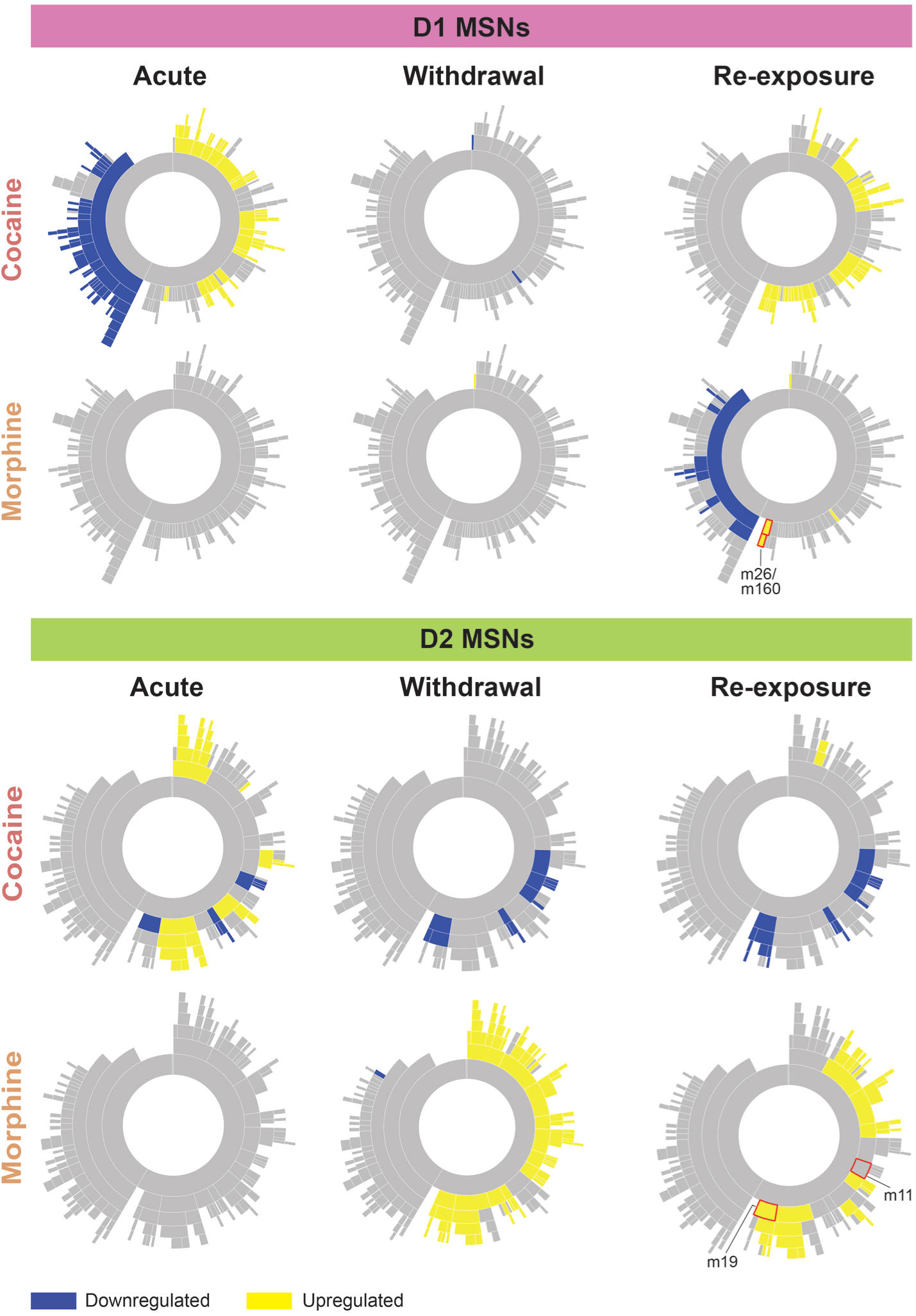
Gene network architecture of cocaine- and morphine-induced transcriptional regulation. MEGENA network for D1 (top) and D2 (bottom) MSNs represented as sunburst plots with segments on each rung representing distinct networks. MEGENA enables progressive network refinement, as reflected in sunburst plots by central rungs being broken into smaller segments moving outwards from central rungs. Note that D1 and D2 MEGENA networks are represented by a single sunburst plot, which is presented repeatedly to highlight condition-specific network enrichment of DEGs. Coloration of sunburst plot segments depicts significant enrichment of upregulated (yellow) or downregulated (blue) genes in particular networks. Modules highlighted with red boxes indicate those emphasized in subsequent figures.

In contrast to cocaine, we found no gene network enrichment from first-ever morphine (SM) or morphine withdrawal (MS). However, re-exposure to morphine after withdrawal (MM) elicited several enriched gene networks. Surprisingly, many of the downregulated genes in MM were also observed in the SC condition, indicating some overlap in gene network regulation in D1 MSNs across cocaine and morphine. It is possible that the gene networks that are enriched in the MM condition exhibit a similar pattern of regulation in SM and MS, but considering their lower absolute magnitude of transcriptional regulation compared to MM (see Figure 1C), the threshold for gene network enrichment was not met.

In D2 MSNs, we found that first-ever cocaine elicits transcriptional responses that enrich both up- and downregulated gene networks, and some of these changes persist through withdrawal and re-exposure. However, we noted that many gene networks engaged by acute cocaine, particularly upregulated networks, lost their enrichment with re-exposure to cocaine after withdrawal. This suggests that, despite the strong overlap in D2 genes activated by cocaine in SC and CC conditions, the CC transcriptional response was consolidated into fewer gene networks. Interestingly, downregulation of several gene networks were conserved across all three cocaine exposure conditions. In contrast to cocaine, first-ever morphine (SM) exposure did not elicit strong enrichment of DEGs across gene networks, which was also observed for D1 MSNs. However, we saw a highly organized transcriptional response, particularly for upregulated genes, through withdrawal from chronic exposure (MS) that was largely preserved with re-exposure after withdrawal (MM). We also found that, across cocaine and morphine, D2 MSNs generally had more overlapping gene networks that were activated across exposure conditions and were conserved across both large and more refined subnetworks. Some gene networks were regulated in opposite directions, while others were consistent across drugs.

Taken together, these results suggest that, as observed in RRHO analysis, chronic cocaine causes a consolidation of transcriptional regulation in D1 MSNs that is preserved at the gene network level. In contrast, chronic morphine appears to progressively amplify gene network regulation in both D1 and D2 MSNs.

### Cocaine and morphine engage overlapping D1 gene networks linked to canonical immediate early gene function

Although we found few gene networks activated by either first-ever or re-exposure to morphine in D1 MSNs, we noted a particular set of modules, m26 and its child module m160 (see Figure 2), that stood out as being commonly activated by re-exposure to cocaine and to morphine after withdrawal (CC and MM). Upon closer examination of the m160 network structure, we found that many canonical IEGs clustered together within these modules (Figure 3A), including *Arc*, *Npas4*, *Egr1*, *Egr3*, *Nr4a3*, and *Fosl2*. As noted above, cocaine exposure caused activation of several IEGs in D1 MSNs, but we were surprised to see that several IEG family members group into gene co-expression networks to potentially influence specific biological processes. Using a nearest neighbor approach, we next isolated subnetworks that interact with IEGs by extracting the two closest partners of each IEG within the broader module 160. This approach found that these IEGs interact with a major hub gene in module 160, which encodes amyloid beta precursor-like protein 2 (*Aplp2*). Notably, APLP2 is critical for neurotransmission, synaptic plasticity, and learning (26), and several indications point to amyloid precursor protein family genes as being involved in CUD and OUD (27,28). Through *Aplp2* and other gene partners in module 160, IEG expression was also linked to the strongest hub gene in module 160, which encodes sodium voltage-gated channel beta subunit 2 (*Scn2b*). SCN2B and plays a critical role in sodium channel function and is necessary for action potential propagation and synaptic calcium dynamics (29).

**Fig 3.**
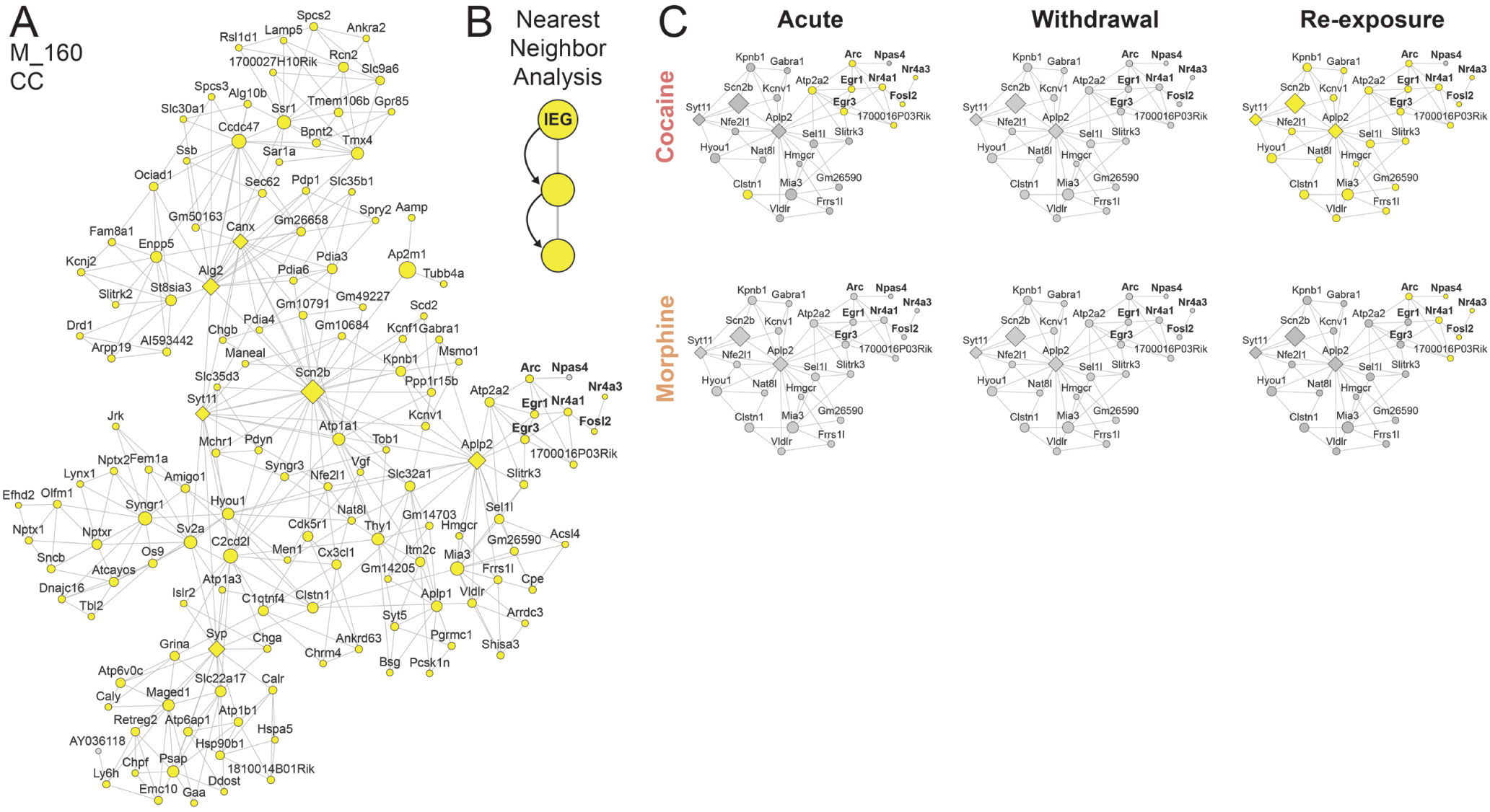
D1 MSN gene network that integrates canonical IEGs is primed for activation by chronic cocaine. **A**, Network structure of module D1 m160 showing defined hubs as diamonds and non-hub nodes as circles, both sized by degree of connections. Immediate early genes are highlighted with bold text. Differentially expressed genes (>20% FC, p<0.05) from the CC condition are overlaid onto this network (upregulated, yellow shading; no change, gray shading), demonstrating near complete activation of this network. **B**, IEG subnetworks were defined by a nearest neighbor approach wherein all nodes 2 steps downstream of IEGs were integrated into a subnetwork module. **C**, IEG subnetworks, which included two influential m160 hub genes, *Aplp2* and *Scn2b*, with overlaid differential gene expression for each experimental condition to show gene network engagement.

We then overlaid differential gene expression results (>20% FC; p<0.05) onto IEG subnetworks in m160 to illustrate how cocaine and morphine exposure influence the expression of these IEGs and how this integrates into shifts in gene network regulation. We found that first-ever cocaine exposure (SC) caused activation of most IEGs in m160, without affecting many of the other genes in the subnetwork. First-ever morphine morphine exposure (SM) had no effect on IEGs or other genes in the subnetwork. Re-exposure to cocaine after withdrawal enabled transcriptional activation to spread from IEGs through hub genes in the network, resulting in a massive wave of transcriptional activation throughout m160, as demonstrated in Figure 3A. These findings support and extend recent work outlining a key role for IEGs in mediating dopamine-dependent D1 MSN responses to acute cocaine (Savell, Day, 2020), and indicate that these genes may integrate into broader networks that influence various biological processes upon repeated drug exposure and withdrawal.

**Fig 4.**
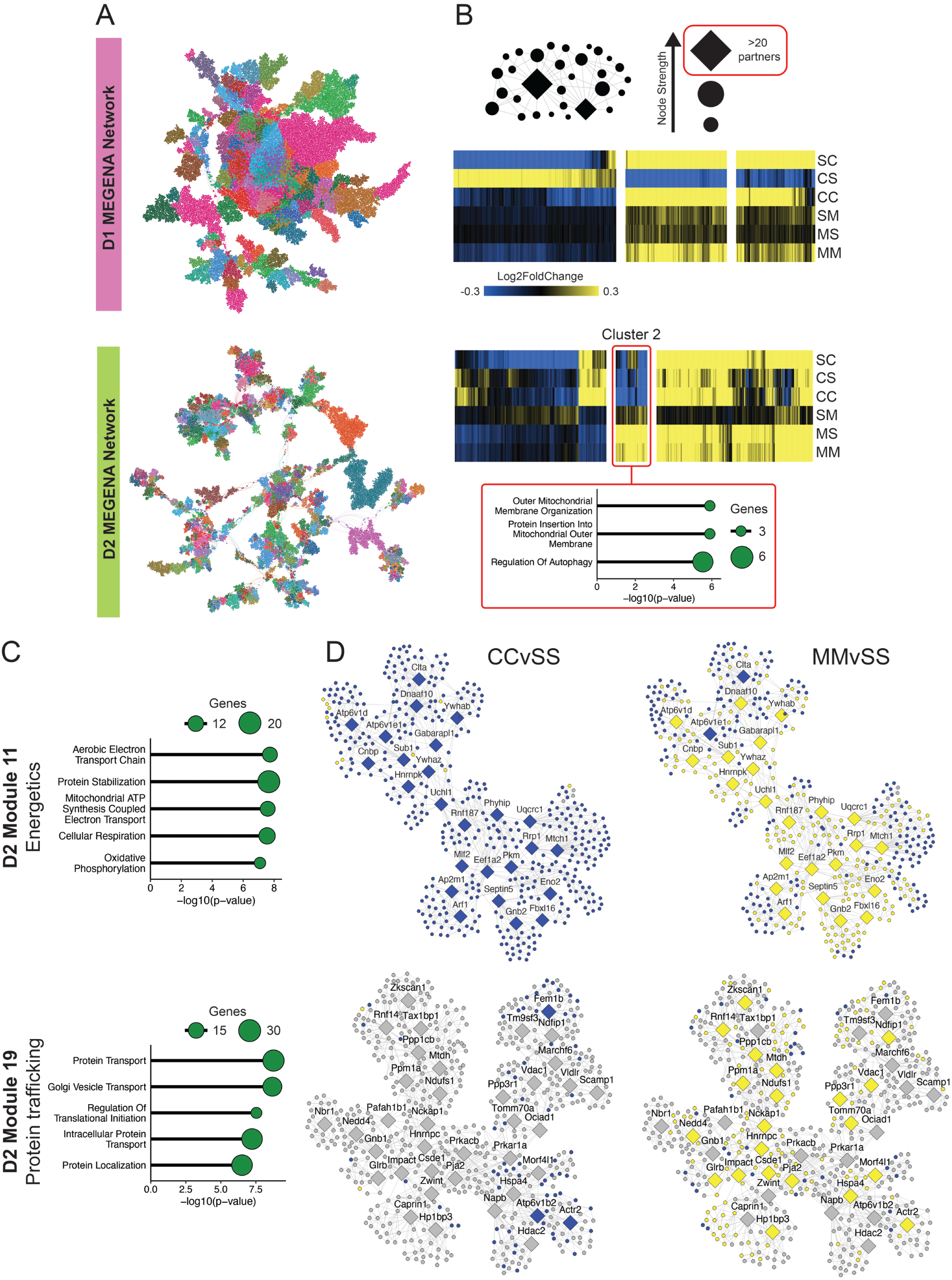
Hub gene analysis reveals key D2 networks that differentiate cocaine- vs. morphine-induced changes to biological functions. **A**, MEGENA network structure for D1 and D2 MSNs. Note, D1 MSNs are more densely interconnected compared to D2 MSNs (mean difference of correlation coefficients p<2.2e-16, see Results). **B**, Hub gene analysis and clustering. Top, scheme describing the approach of extracting top-ranked gene network hubs for D1 or D2 MSNs based on exhibiting more than 20 co-expression partners. Heatmaps show expression patterns in log2FoldChange (up, yellow; down, blue) for each top-ranked hub gene across experimental conditions after hierarchical clustering. Bottom chart shows results of gene ontology analysis of biological processes enriched in D2 cluster 2 which exhibits opposite regulation between cocaine and morphine exposure conditions. **C**, Gene ontology analysis of two D2 MEGENA modules with most represented in D2 cluster 2, showing enrichment of biological processes associated with energetic utilization (m11) and protein trafficking (m19). **D**, Gene network structure of m11 and m19 presenting hub genes as diamonds and non-hub nodes as circles. Differentially expressed genes (p<0.05) from CC (left) and MM (right) conditions are overlaid showing upregulated (yellow) or downregulated (blue) genes, throughout the network.

Interestingly, while re-exposure to morphine post-withdrawal activated all IEGs, this activation did not mirror the hub gene activation within the subnetwork, akin to the patterns seen with the initial cocaine exposure. Notably, withdrawal from cocaine or morphine had no effect on the m160 subnetwork or enrichment of DEGs as a whole, indicating that the changes observed for both drugs require pharmacological induction. This observation indicates that the recurrent activation of this IEG cluster might facilitate extensive transcriptional shifts in the network, particularly through the pivotal hub genes *Aplp2* and *Scn2b*. This result suggests that chronic cocaine, and potentially morphine, drives gene priming mechanisms that may precipitate pronounced biological responses upon re-exposure following protracted abstinence.

### Patterns of gene network consolidation of cocaine and morphine responses across D1 and D2 MSNs

Our transcriptome-wide RRHO results (Figure 1B) indicated that D1 and D2 MSNs exhibit a variety of transcriptional patterning in response to cocaine and morphine that spanned acute exposure, withdrawal, and re-exposure conditions. Based on these findings, we determined whether these patterns of gene regulation were maintained and consolidated within gene network architecture, subserving particular biological functions. To narrow the dimensionality of the overall D1 and D2 MSN gene networks (Figure 4A), we ranked all nodes in each network by their relative strength of association with other nodes and selected all genes that exhibit >20 bona fide connections to generate a list of the top-ranked hubs in each network (Figure 4B). This analysis yielded 369 hub genes for D1 MSNs and 365 hub genes for D2 MSNs. Specifying hub genes enabled us to examine patterns of transcriptional regulation of highly influential genes that control global expression throughout broader gene networks. We then performed hierarchical clustering analysis to survey patterns of gene regulation across experimental conditions, as represented by heatmaps in Figure 4B.

In D1 MSNs, we found that acute cocaine exposure (SC) produced strong transcriptional responses in hub genes that were largely recapitulated by re-exposure to cocaine after withdrawal (CC). However, this re-exposure appeared to drive consolidation of hub gene regulation based on a diminished transcriptional response compared to SC. This cocaine response trajectory was largely paralleled in D2 MSNs. We also noted that in D1 MSNs, as observed in RRHOs, withdrawal from cocaine (CS) produced opposite patterns of hub gene regulation compared to either cocaine exposure condition (SC or CC) – a phenomenon absent in D2 MSNs. In contrast, acute morphine exposure (SM) induced only subtle transcriptional shifts in both cell types, but these changes became more pronounced with prolonged morphine exposure (both MS and MM). Notably, extended withdrawal from chronic morphine (MS) triggered a robust transcriptional shift in D2 MSNs, echoing the changes seen upon re-exposure (MM), an effect that was not observed in D1 MSNs.

### D2 MSN gene networks associated with key biological processes distinguish molecular signatures of cocaine vs. morphine exposure

Transcriptome-wide RRHO analyses (Figure 1B) revealed that D2 MSNs exhibit a particularly unique transcriptional response to cocaine and morphine across acute exposure, withdrawal, and re-exposure phases. Additionally, clustering analysis of hub genes found generally less overlap across cocaine and morphine exposure conditions in D2 MSNs relative to D1 MSNs. Thus, we wondered whether unique transcriptional regulatory mechanisms that separate cocaine vs. morphine responses in D2 MSNs could be classified within defined gene networks. Our hierarchical clustering analyses pinpointed a set of hub genes (cluster 2) in D2 MSNs that were consistently suppressed across all cocaine exposure scenarios but upregulated in all morphine conditions (Figure 4B). Gene ontology analysis of the genes in this cluster revealed an enrichment of processes related to mitochondrial structure and function as well as autophagy.

To discern if these drug-specific alterations in hub gene expression permeated broader gene networks and, in turn, influenced distinct cellular functions, we turned our attention to the most prominent D2 MSN MEGENA networks associated with this hub gene cluster: m11 and m19 (Figure 4C; see sunburst plots in Figure 2). Of the total 34 hub genes, 11 belonged to m11, representing 44% of MEGENA-defined hub nodes in this module, while 14 belonged to m19, representing 36% of this module’s hub nodes. Gene ontology analysis of these modules, independent of DEG expression, revealed that module 11 was associated with biological processes related to energetic utilization and oxidative phosphorylation, while module 19 is associated with biological processes related to protein trafficking and Golgi function. We then overlaid differentially expressed genes (nominal p<0.05) from the MM and CC conditions to examine how gene priming or desensitization after withdrawal from chronic drug exposure distributes throughout these two modules (Figure 4D). In m11, the CC condition elicits broad downregulation of nearly every gene in this network, while in the MM condition nearly all hub genes are upregulated in addition to the majority of genes immediately connected to the hub genes. Interestingly, however, many nodes on the outskirts of the module are also downregulated, suggesting that this network, although divergent between cocaine and morphine, may exhibit particularly dynamic regulation. In m19, cocaine re-exposure drives only minimal gene network regulation that is largely restricted to a handful of downregulated genes and hubs. Morphine re-exposure, on the other hand, elicits broad activation of this gene network. These results suggest that key gene networks associated with active cellular processes that require protein mobilization and increased mitochondrial function for intracellular processes are oppositely regulated in NAc D2 MSNs by cocaine vs. morphine. These core networks may serve as critical divergent factors that support separable functional and behavioral sequelae of cocaine and morphine action, especially during periods of abstinence and relapse.

In our hierarchical clustering analysis, we noted another group of hub genes that exhibited opposite regulation between cocaine and morphine conditions, but in this case they were upregulated by cocaine and mostly downregulated by morphine (Supp Fig 2A). Using gene ontology analysis, we found that this hub gene cluster enriched for biological processes related to extracellular matrix (ECM) maintenance and associated collagen biology. This finding was particularly interesting given our recent work profiling transcriptional regulation in a mouse model of OUD, as well as recent findings in postmortem NAc from patients with OUD, both of which implicate an important role for ECM dysregulation in supporting opioid addiction pathology (11,12,30). Nearly all genes in this cluster were associated with module 3 (see sunburst plots in Figure 2), a large gene network containing nearly 5700 genes (Supp Fig 2B). Gene ontology analysis of this large module revealed ECM functions as the top enriched biological process in this network. Overlaying differentially expressed genes (p<0.05) onto module 3 (Supp Fig 2C), we found that acute cocaine (SC) produces modest changes with both up- and down-regulation throughout the network, but cocaine withdrawal (CS) produces largely upregulated genes which is further exacerbated by cocaine re-exposure (CC). On the other hand, acute morphine (SM) produces modest downregulation of genes in m3, while morphine withdrawal (MS) causes profound suppression of gene expression distributed throughout the gene network, which is maintained when re-exposed to morphine (MM) but blunted compared to withdrawal. While morphine acts on genes throughout this network, cocaine responses were largely conserved to a subnetwork of genes (Supp Fig 2D), which also enriched for processes related to extracellular matrix function. Taken together, these results support the idea that cocaine and morphine drive separable responses in D2 neurons, with some of these changes relating to divergent extracellular matrix biology to support opposite functional changes.

## Discussion

Cocaine and morphine, despite their shared ability to induce addiction-related behavioral abnormalities, drive partly distinct molecular responses in the NAc, a brain region crucial for reward processing and motivation. Here, we provide a comprehensive overview of cell-type-specific transcriptional regulation in the NAc across multiple phases of abuse-relevant exposure to cocaine and morphine. Our findings reveal intricate patterns of D1 and D2 MSN gene regulation across different stages of drug exposure, withdrawal, and relapse.

Cocaine’s influence on D1 MSNs is characterized by dynamic transcriptional responses, with an inversion of transcriptional regulation observed during prolonged withdrawal—a phenomenon not mirrored by morphine. These findings suggest that the molecular underpinnings of cocaine’s acute effects on D1 MSNs and its stimulus-responsive gene networks are unique. Interestingly, gene networks integrating IEGs appear to be particularly sensitive to increasing drug exposure, irrespective of the drug class. While cocaine acts more potently and directly on IEGs as potential gateways into broader gene networks, morphine also exerts an increasing influence on the network following withdrawal, potentially amplified by repeated exposure. Interestingly, one key hub gene in this network that served as a direct coexpression partner with several IEGs was *Aplp2*. Notably, *Aplp2*, which was also significantly upregulated in the NAc of OUD patients (12), and its closely related genes *Aplp1* and *App*, have been linked to both CUD and OUD in humans (27,28). Given that IEGs respond to strong stimuli, it is plausible that a threshold of concerted IEG activation enables increasingly broad gene network activity. This engagement might become primed with a history of drug exposure, facilitating rapid and broad gene regulation upon re-exposure, as observed for cocaine. The absence of changes during withdrawal, when no drug is present, further supports the idea that these gene networks and IEGs are activated in response to a proximal stimulus. The overlap in overall transcriptional responses in D1 MSNs induced by cocaine and by morphine suggests a shared mechanism, emphasizing neuroplastic changes common to both drugs. Such a shared molecular mechanism could be an ideal target for therapeutic interventions.

The molecular landscape of D2 MSNs starkly segregates the effects of cocaine and morphine. While D1 MSNs exhibit significant overlap in transcriptional responses when drug is present, D2 MSNs show largely no overlap or opposite patterns of regulation across drug classes. Many of the patterns of broad transcriptional regulation were maintained at the gene network level, particularly in terms of hub gene expression. We found that several D2 gene networks exhibited completely opposite engagement by cocaine and morphine. The ability of hub genes to influence broad waves of transcriptional regulation suggests that even minor perturbations at the hub level can drive extensive transcriptomic changes, potentially explaining the divergent gene regulation observed between cocaine and morphine in D2 MSNs. We found that these oppositely regulated networks coordinated crucial biological processes associated with mitochondrial functions, protein trafficking, and ECM maintenance, in addition to many others. These findings represent points of cell-type-specific disruptions that may segregate functional and behavioral responses to psychostimulants and opioids, providing novel interventional targets for CUD and OUD.

Results from withdrawal conditions emphasize that D1 MSNs coordinate highly dynamic transcriptional regulation in response to drugs of abuse, while D2 MSNs appear to accumulate changes associated with chronic exposure. Despite opposite regulation patterns, D2 MSNs exhibit increasingly strong, parallel magnitudes of gene expression with increasing exposure to both cocaine and morphine, while D1 MSNs largely showed a diminished or opposite regulation. Transcriptional regulation that persists through withdrawal in D2 MSNs was also maintained into organized gene networks. In contrast, almost no gene network enrichment is observed for D1 MSNs through withdrawal. Morphine exemplifies this showing a particularly strong transcriptional response in many of the same genes activated by re-exposure that were maintained and expressed at higher magnitudes through withdrawal. Stronger transcriptional responses in withdrawal may relate to processes associated with the idea of D2 “super-sensitivity” with protracted opioid abstinence (31), and our gene network analysis points to key molecular pathways that may mediate these unique effects of opioids.

Our findings underscore the importance of tailoring treatment strategies across SUDs. The clinical manifestations of CUD and OUD, while sharing similarities, also exhibit distinct characteristics. Our results suggest that some of the divergent aspects of CUD and OUD may relate to unique effects of the two drug classes on two prominent cell types within the NAc, a crucial hub in the reward system. The unique influences of cocaine and morphine on influential gene network hubs suggest that, despite broad transcriptional overlap, many gene networks likely involved in key biological functions exhibit drug-divergent responses. This divergence, while expected given the known functional and molecular differences between cocaine and morphine, emphasizes the need for more effective, tailored treatment strategies for CUD and for OUD. These results also highlight the importance of surveying cell-type-specific molecular regulation to comprehensively determine molecular mechanisms governing addiction pathology.

In conclusion, these studies shed light on the intricate molecular landscapes of D1 and D2 MSNs in response to cocaine and morphine exposure. The distinct transcriptional patterns, and their implications for the potentially unique molecular mechanisms involved, underscore the need for a deeper understanding, which could pave the way for more effective, tailored therapeutic strategies for SUDs.

## Methods

### Animals

Male double-transgenic mice expressing D1- or D2-specific nuclear-tagged GFP were generated by crossing LSL-eGFP::L10a (IMSR_JAX:022367) mice with either D1-Cre (MGI:3836633) or D2-Cre (MGI:3836635) mice. All mice were bred in-house on a C57BL6/J background and were 8-20 weeks old at the beginning of experimental procedures. Mice were housed on a regular light-dark cycle with lights ON at 7:00 am, and food and water available ad libitum for the duration of the study. All experiments were conducted in accordance with the guidelines of the Institutional Animal Care and Use Committee (IACUC) at Mount Sinai (protocol number 08-0465).

### Drug treatment

Cocaine hydrochloride (from the National Institute on Drug Abuse) and morphine sulphate (from the National Institute on Drug Abuse) were dissolved in 0.9% saline and administered intraperitoneally at doses of 20 mg/kg or 10 mg/kg, respectively. These doses were chosen based on their roughly equivalent rewarding effects, as established in prior conditioned place preference experiments (21). In the cocaine cohort, mice received saline or cocaine each day for 10 days prior to withdrawal, while the morphine cohort received 15 treatments. As depicted in Figure 1A, after 30 days of homecage withdrawal, mice received challenge injections of saline, cocaine, or morphine and were sacrificed 1h later.

### Tissue processing and fluorescence activated nuclei sorting (FANS)

Mice were euthanized by cervical dislocation, and bilateral NAc tissue was rapidly dissected on ice from 1-mm thick coronal sections using a 14G punch and flash frozen on dry ice. Nuclei were purified as previously described (21). Frozen NAc samples were first homogenized in lysis buffer (0.32 M sucrose, 5 mM CaCl2, 3 mM Mg(Ace)_2_, 0.1 mM EDTA, 10 mM Tris-HCl) using high-clearance followed by low-clearance glass douncers (Kimble Kontes). Homogenates were then passed through a 40um cell strainer (Pulriselect) into ultracentrifuge tubes (Beckman Coulter). A 5ml high-sucrose cushion (1.8 M sucrose, 3 mM Mg(Ace)2, 1 mM DTT, 10 mM Tris-HCl) was then underlaid, and samples were centrifuged at 24,000 rpm at 4°C for 1h in a SW41Ti Swinging-Bucket Rotor (Beckman Coulter). After discarding the supernatant, nuclei pellets were resuspended in 500ul of ice-cold phosphate-buffered saline, and DAPI (1:5000) was added. All solutions contained inhibitors for RNAse (SUPERase-in, Invitrogen) and ribonuclease (RNasin Recombinant, Promega) at concentrations of 1:1000 (sucrose buffer) or 1:500 (lysis buffer and PBS). Nuclei were then immediately sorted using a BD FACS Aria II three-laser system with a 100um nozzle. Debris and doublets were first gated based on forward scatter and side scatter, nuclei were then gated as DAPI-positive events (Violet1-A laser), and GFP positive events were sorted directly into 1.5 ml Eppendorf tubes containing TriZol LS (Ambion) and flash frozen on dry ice. Samples ranged from 18,000-60,000 GFP+ nuclei collected for downstream analysis.

### RNA sequencing

After FANS, nuclei were thawed on ice and homogenized in TriZol, and RNA was extracted using the Direct-zol RNA microprep kit (Zymo Research) according to manufacturer instructions. To generate sequencing libraries, 1ng of total RNA was used as input for Takara SMARTer® Stranded Total RNA-Seq Kit v3 - Pico Input Mammalian kit, with ribodepletion, according to manufacturer’s instructions. Sequencing libraries were generated for each sample individually using Takara’s Unique Dual Index Kit. Following library preparation, sequencing was performed with Genewiz/Azenta on an Illumina Novaseq S4 machine with a 2x150 bp paired-end read configuration produce 40M reads per sample. Samples from cocaine and morphine cohorts were sequenced separately. Within cocaine or morphine cohorts, all samples were multiplexed and run concurrently. Quality control was performed using FastQC (www.bioinformatics.babraham.ac.uk/projects/fastqc/). All raw sequencing reads underwent adapter trimming using Trimmomatic (github.com/usadellab/Trimmomatic) and were mapped to mm10 using HISAT2 (daehwankimlab.github.io/hisat2/). Duplicate reads were removed using Picard MarkDuplicate tool (broadinstitute.github.io/picard/), and count matrices were generated using the featureCounts function of the subread package (subread.sourceforge.net/featureCounts.ht). Normalized counts were filtered to include only the top half of the genome, yielding a minimum base mean of ∼24 for D1 cocaine, ∼15 for D2 cocaine, ∼33 for D1 morphine, and ∼33 for D2 morphine. Differential expression was performed on this filtered gene list in R version 4.0.2 using the DESeq2 package version 1.28.1 (32), with built-in independent filtering disabled. Experimental conditions were always compared to their respective control group (i.e. SC vs. SS, CS vs. SS, CC vs SS, etc.). Significance for DEGs was set at either 20% expression fold change (FC ± 20%) and p<0.05, and p<0.05 only as specified throughout.

### Rank-rank hypergeometric overlap

RRHO enables comparison of gene expression profiles across experimental conditions in a threshold-free manner (33). RRHO2 plots, which illustrate both concordant and discordant genes (34) were generated comparing differential gene expression lists from cocaine versus morphine conditions (e.g. SC v SM) or across cell types (eg. D1 and D2 MSNs for the SC condition). RRHO2 plots were generated in R using the RRHO2 package with default settings (github.com/RRHO2/RRHO2).

### Heatmaps

Union heatmaps were generated from a master list of all differentially expressed genes (>20% FC, p<0.05) identified across experimental conditions with duplicates removed. Raw log2FoldChange values for all genes in this master list were then plotted for all experimental conditions using Morpheus (https://software.broadinstitute.org/morpheus). Hierarchical clustering (Figure 4) was performed using Morpheus, according to Euclidean distance (with Linking method set to Average), and dendrograms were split at the second branch to visualize the top 3 clusters identified.

### Gene ontology (GO) analysis

Enrichr (35) was used to query the 2023 GO:BP database of the Gene Ontology Consortium (Gene Ontology Consortium, 2021) via the EnrichR package in R (github.com/wjawaid/enrichR). Data was visualized in R using the tidyverse package, version 1.3.1.

### Multiscale embedded gene co-expression network analysis (MEGENA)

MEGENA was performed as described previously (24). Briefly, gene co-expression networks were constructed using the R package MEGENA (github.com/songw01/MEGENA) collapsing all experimental conditions from both cocaine (SS, SC, CS, CC) and morphine (SS, SM, MS, MM) cohorts into a single network representing D1 or D2 MSNs. Samples with less than 10 million reads were excluded from MEGENA network construction, yielding a sample size of 55 for the D1 MSN network and 50 for the D2 MSN network. The D1 network had 41,095 edges, and the absolute values of the correlation coefficient ranged from 0.911912 to 1 (mean value 0.96824, SD = 0.019221), while the D2 network had 38,724 edges, and the absolute values of the correlation coefficient ranged from 0.860763 to 1 (mean value 0.94304, SD = 0.034624). Modules with less than 50 genes were excluded from analysis. Hub genes for each module were determined using a combination of node strength and node degree with a 5% false discovery rate (FDR). Enrichment p-values were calculated using the Fisher’s exact test and adjusted for multiple comparisons using the Benjamini-Hochberg method. Sunburst plots depicting network organization and DEG enrichment were generated using the R package sunburstR (github.com/timelyportfolio/sunburstR).

### Gene network visualization

MEGENA network architecture was visualized using Cytoscape version 3.10.0 (36). Overall D1 or D2 gene networks were built using connection weight to define node associations. Node degree of connectivity was then included to establish node size scaling. Overall networks were then filtered into modules of interest, and module-specific hub genes were presented as diamonds while non-hub genes were represented with circles. Subnetworks were then organized using a prefuse force directed layout followed by manual rearrangement to shorten path lengths. Differential gene expression values (p<0.05) were then included for each node to illustrate upregulation (yellow) or downregulation (blue) of network components. Immediate early gene subnetworks presented in Figure 3 were generated by selecting all primary and secondary nodes using the “First neighbor of selected node” tool after selecting all IEGs.

## Acknowledgements

We thank Stephen Pirpinias, Katherine Beach, Catherine McManus, Kyra Schmidt, and Ezekiell Mouzon for transgenic breeding and genotyping. We also thank Dr. Edgardo Aritzia from the Dean’s Flow Cytometry CoRE at the Icahn School of Medicine at Mount Sinai for assistance in nuclei sorting. This work was supported by National Science and Research Council of Canada (NSERC PDF to C.J.B.), a National Institute on Alcohol Abuse and Alcoholism (K99AA027839 to P.M.), from the Brain and Behavior Research Foundation (BBRF Young Investigator Grant to P.M.) and the National Institute of Health (NIDA P01DA047233 to A.S., P.J.K., Y.L.H., L.S., and E.J.N).

## Author Contributions

Conceptualization: C.J.B., P.M., P.J.K., and E.J.N. Methodology: X.Z., A.S., Y.S.L., and B.Z. Investigation: C.J.B., P.M., L.M.H., X.Z., R.F., M.E. Visualization: C.J.B., P.M., X.Z., Formal analysis: C.J.B., P.M., X.Z., M.E. Writing: C.J.B., P.M., E.J.N. Writing–review and editing: All authors read and commented on the manuscript. Supervision: A.S, P.J.K., L.S., B.Z., E.J.N.

## Data and materials availability

All data needed to evaluate the conclusions in the paper are present in the paper and/or the Supplementary Materials. All RNAseq data reported in this study will be deposited publicly in the Gene Expression Omnibus upon manuscript acceptance. Other supporting scripts/code used in this study are available from the corresponding author upon request.

---

**Supp Fig 1.**
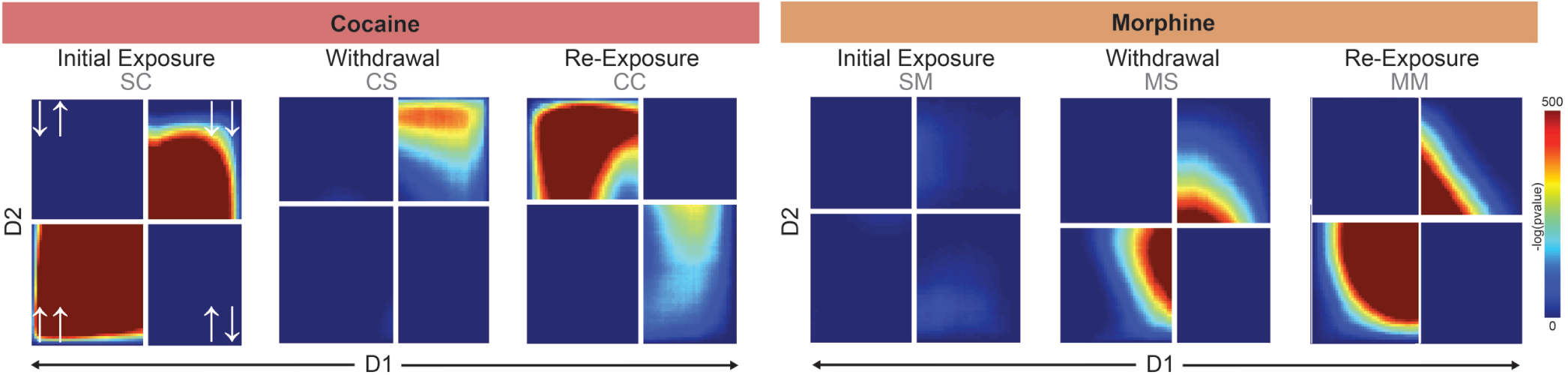
Patterns of transcriptome-wide regulation across D1 and D2 MSNs induced by cocaine and morphine. Rank-rank hypergeometric overlap plots compare threshold-free gene regulation between D1 and D2 MSNs for first-ever drug exposure, withdrawal, or re-exposure after withdrawal. Heat indicates strength of overlap, and quadrants represent direction of gene expression (white arrows; lower-left quadrant, genes up in both; upper-right quadrant, genes down in both; upper left quadrant, genes down in D2 but up in D1; lower right quadrant: genes up in D2 but down in D1).

**Supp Fig 2.**
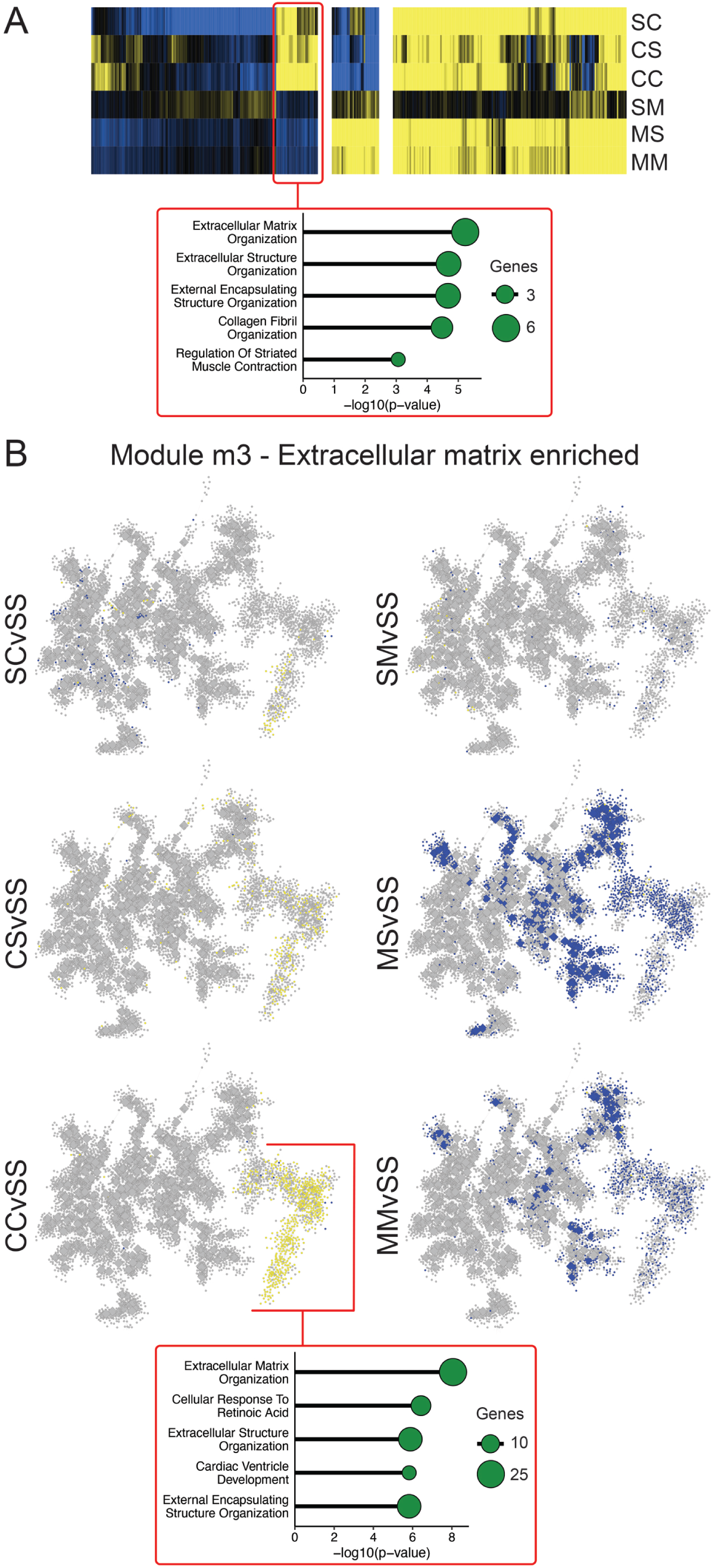
Cocaine and morphine oppositely regulate a D2 gene network coordinating ECM dynamics. **A**, Hierarchical clustering of D2 MEGENA network hub genes from Figure 4, highlighting a group of hub genes showing upregulation (yellow) in cocaine conditions, but downregulation (blue) in morphine conditions. Bottom of panel shows gene ontology analysis of this cluster significantly enriching for biological processes associated with extracellular matrix-associated functions. **B**, Network structure of D2 module m3, a large module of nearly 5700 genes from which nearly all hub genes in the cluster analysis originated. Gene ontology analysis found extracellular matrix organization to be the top enriched biological process associated with m3 (Benjamini-Hochberg corrected p<0.001). Differential expression results (p<0.05) from all experimental conditions are overlaid onto m3, demonstrating a near complete divergence between cocaine, which upregulates (yellow) many genes in the network, and morphine, which downregulates (blue) many genes in this network. Notably, cocaine effects (particularly CC) appeared to mostly cluster in a subnetwork within m3, which also enriched specifically for extracellular matrix function, and are largely downregulated by morphine (particularly MM).

